# Diffuse phenotypic structure in native chickens across Java and Wallacea is consistent with human-mediated connectivity and decentralized selection

**DOI:** 10.64898/2026.09.08.750014

**Authors:** N. Widyas, S. Fiddaman, S. Prastowo, A. Ratriyanto, L. A. F. Frantz, R. Drinkwater, R. Kusumaningrum, A. E. T. Sulfiar, E. Wera, P. M. Bulu, Sulasmi, P. R. Matitaputty, T. S. S. Panjaitan, R. Widyastuti, A. Setiaji, A. B. Dharmayanthi, A. Pratiwi, L. A. Pradista, T. Nugroho, Z. A. Wahid, G. Pambuko, E. D. Novitasari, A. D. Sayogo, A. L. Smith

## Abstract

**Objective:** This study aimed to investigate morphometric and plumage-colour variation among native chickens from Wallacea and Java, Indonesia, and to evaluate the contribution of geographic and production-system factors to phenotypic diversity.

**Methods:** Native chickens were sampled from 259 households across eight provinces in Indonesia. Household-level information on production systems and farmer selection practices was collected through structured surveys. Morphometric traits and HSV-derived plumage-colour variables were analysed using principal component analysis (PCA), permutational multivariate analysis of variance (PERMANOVA), and hierarchical variance partitioning to assess phenotypic variation across locations, provinces, and islands.

**Results:** Substantial phenotypic variation was observed among sampled populations, with significant province-and island-level differences detected for several morphometric traits. However, extensive overlap among locations was evident in both morphometric and plumage-colour space, indicating diffuse rather than discrete structuring. Within-location variation accounted for the largest proportion of total phenotypic variance across most traits, consistent with decentralized smallholder management and continued human-mediated exchange.

**Conclusion:** Geographic fragmentation and agroecological heterogeneity alone appear insufficient to generate strongly differentiated phenotypic groups when connectivity among populations is maintained. Whether this phenotypic connectivity corresponds to contemporary gene flow, retained ancestral variation, or convergent environmental responses remains to be tested using genomic data.

**Implications:** This study shows that native chickens in Indonesia maintain high diversity despite being raised across different islands and environments. The findings support protecting local chicken resources that underpin smallholder livelihoods, rural food security, and cultural traditions, and can help future breeding and conservation programs develop chickens better adapted to local climates, diseases, and farming conditions: improving farmer income and sustainable local poultry production.

## Introduction

Genetic and phenotypic variation within and among populations reflects the interplay between gene flow, genetic drift and selection: restricted dispersal and geographic barriers tend to promote differentiation among populations, whereas high connectivity and weak selection maintain variation within populations and reduce spatial structure [1,2,3,4].

While these principles are well established for natural populations, their manifestation in domesticated species is more complex: domestication is increasingly recognised as a dynamic, reticulate process involving multiple origins, recurrent admixture, and continuous gene flow between populations [5,6]. In chickens, genetic evidence indicates contributions from multiple junglefowl species and extensive introgression, producing heterogeneous genetic backgrounds rather than discrete lineages [7,8], challenging simple expectations of population structure based on geographic isolation.

The complexity of domesticated systems is further amplified in smallholder production environments. Village farming systems for native chickens, widely distributed across Africa and Asia, are characterised by small flock sizes, low-input management, and the absence of formal breeding programmes [9,10], with animals frequently exchanged among households, largely uncontrolled mating, and decentralised selection based on observable traits rather than systematic performance recording [11,12]. Under these conditions, farming systems may themselves shape the evolutionary processes contributing to genetic and phenotypic variation, examined here indirectly through morphometric, plumage-colour, and production-system data rather than direct genetic measurement.

Empirical studies of native chickens consistently report high genetic and phenotypic variation within populations and limited differentiation among them [13,14,15], interpreted through interacting processes including gene flow, genetic drift, and heterogeneous selection rather than strong directional selection or long-term isolation. A key gap remains, however, in understanding how these processes interact across spatial scales and environmental contexts in regions with complex biogeographic histories.

Indonesia provides a unique setting to address this question. The Wallacea region, between the Asian and Australasian faunal zones, has long been recognised as a natural laboratory for studying evolutionary processes, where geographic barriers such as the Wallace Line contribute to pronounced differentiation in wild species [16,17]. Whether such biogeographic patterns are retained, modified, or overridden by human-mediated processes in domesticated species remains unclear; previous genetic studies of Indonesian chickens reveal substantial diversity and admixture, suggesting variation reflects both historical and contemporary processes [18,19]. Although Indonesian native chickens are phenotypically diverse and are managed under heterogeneous smallholder systems, it remains unclear how their phenotypic variation is partitioned among islands, provinces, sampling locations, and individuals within locations. Moreover, few studies have jointly examined this hierarchical phenotypic structure and the household practices that may contribute to its maintenance.

Addressing these questions requires integrating phenotypic data with information on farming systems and spatial structure. In smallholder livestock systems, spatial phenotypic structure may arise from geographic separation, environmental heterogeneity, farmer-mediated selection, and human-mediated animal movement: frequent exchange among households can maintain connectivity and potentially facilitate gene flow, while decentralized, heterogeneous selection may limit consistent directional differentiation. However, phenotypic data alone cannot distinguish contemporary gene flow from retained ancestral variation or convergent environmental responses, so phenotypic patterns should be interpreted together with animal-exchange and farmer-management information, while direct inference of genetic connectivity requires genomic data.

In this study, we investigate the structuring of phenotypic diversity in Indonesian local chickens across Java and multiple islands within the Wallacea region. Using morphometric and plumage-colour traits, combined with farming-system and selection-practice information, we examine how phenotypic variation is distributed across hierarchical spatial levels, among regions, among locations, or within locations, providing a system-level framework for understanding how animal genetic resources are generated, maintained, and structured in decentralized livestock systems.The novelty of this study lies in integrating quantitative morphometric and plumage-colour measurements with household-level management and selection information across hierarchical geographic scales.

## Materials and Methods

### Ethics approval

All procedures involving animals were approved by the Animal Ethics Committee, Faculty of Animal Science, Universitas Sebelas Maret (Approval No. 150.2/UN27.14/TA.00.03/2026, dated 6 February 2026), and were conducted in accordance with relevant institutional and national guidelines for the care and use of animals. The study was observational and involved non-invasive measurements of animals under field conditions.

### Study area and sampling design

The study was conducted across three provinces in Java (Central Java, East Java, and West Java) and five provinces within the Wallacea region (Maluku, North Maluku, Southeast Sulawesi, West Nusa Tenggara, and East Nusa Tenggara), which differ in biogeographic history, agroecological conditions, altitude, climate, and market accessibility, providing a suitable framework for examining phenotypic variation across heterogeneous smallholder environments.

Sampling locations were selected to represent typical smallholder chicken production systems. Within each location, households keeping local chickens were identified through local networks, and birds were sampled opportunistically to capture existing variation under village conditions. A total of 612 chickens from 259 households were included (Table 1), covering 13 island systems (Java, Lombok, Sumbawa, Timor, Flores, Sumba, Ambon, Seram, Halmahera, Ternate, Tidore, Buton, and Muna) spanning conditions from humid highland systems in Java to seasonally dry and humid island environments in eastern Indonesia (Table 2). Only apparently adult female chickens were sampled, since females represent the reproductive core of village flocks under continuous farmer-driven selection, whereas males may be disproportionately affected by sex-limited ornamental traits and non-random trading, reducing confounding from sex-specific traits.

**Table 1.** Distribution of sampled households and chickens across locations.

| Region | Location | Number of respondents | Number of chickens sampled | Average flock size |
| --- | --- | --- | --- | --- |
| Wallacea | Maluku | 42 | 100 | 23.93 ± 23.53 |
| Wallacea | North Maluku | 47 | 100 | 22.02 ± 18.44 |
| Wallacea | Southeast Sulawesi | 28 | 74 | 14.89 ± 8.85 |
| Wallacea | East Nusa Tenggara | 50 | 92 | 16.02 ± 10.95 |
| Wallacea | West Nusa Tenggara | 54 | 107 | 12.76 ± 14.21 |
| Java | Central Java | 16 | 40 | 17.25 ± 14.48 |
| Java | East Java | 12 | 36 | 18.75 ± 13.55 |
| Java | West Java | 10 | 63 | 14.60 ± 7.00 |

**Table 2.** Agroecological characterization of native chicken sampling sites.

| Province | Location | Altitude range (m a.s.l.) | Mean annual temp (°C) | Rainfall type | Agroecological characterization | Island system |
| --- | --- | --- | --- | --- | --- | --- |
| West Java | Jatinangor | 700–900 | 22–24 | High rainfall | Humid highland | Java |
| West Java | Sumedang | 400–800 | 22–24 | High rainfall | Humid highland | Java |
| Central Java | Temanggung | 600–1,200 | 20–22 | High rainfall | Volcanic highland | Java |
| East Java | Magetan | 500–900 | 22–24 | Seasonal–high rainfall | Upland volcanic | Java |
| West Nusa Tenggara | Batukliang | 50–200 | 26–28 | Seasonal rainfall | Dry lowland | Lombok |
| West Nusa Tenggara | Bayan | 100–400 | 25–27 | Seasonal rainfall | Dry upland | Lombok |
| West Nusa Tenggara | West Sumbawa | 0–200 | 27–29 | Seasonal rainfall | Dry coastal | Sumbawa |
| East Nusa Tenggara | Kupang | 0–200 | 27–29 | Low seasonal rainfall | Dry lowland | Timor |
| East Nusa Tenggara | Manggarai | 200–1,200 | 24–26 | Seasonal rainfall | Mountainous tropical | Flores |
| East Nusa Tenggara | Southwest Sumba | 0–300 | 26–28 | Low seasonal rainfall | Savanna lowland | Sumba |
| Maluku | Ambon | 0–200 | 26–28 | High rainfall | Humid coastal | Ambon |
| Maluku | Central Maluku | 0–100 | 26–28 | High rainfall | Humid island | Seram |
| Maluku | West Seram | 0–200 | 26–28 | High rainfall | Humid forest–coastal | Seram |
| North Maluku | Ternate | 0–200 | 26–28 | High rainfall | Volcanic island | Ternate |
| North Maluku | Tidore | 0–200 | 26–28 | High rainfall | Volcanic island | Tidore |
| North Maluku | Sofifi | 0–50 | 27–29 | High rainfall | Coastal humid | Halmahera |
| North Maluku | West Halmahera | 0–300 | 26–28 | High rainfall | Humid forested island | Halmahera |
| Southeast Sulawesi | Buton | 0–300 | 27–29 | Seasonal rainfall | Dry coastal island | Buton |
| Southeast Sulawesi | Muna | 0–200 | 27–29 | Seasonal rainfall | Dry island lowland | Muna |

### Production system characterization

Information on farming systems and management practices was collected through structured interviews with chicken-keeping households, recording flock size, rearing system, housing, feeding practices, access to forage, sources of replacement animals, and the frequency and mode of animal exchange (purchase, sale, and informal transfer).

Farmer selection practices were documented by recording criteria used for retaining breeding animals and for culling, including whether decisions were based on observable traits (e.g., body size, plumage) or performance-related attributes. These data characterized the production system as a set of conditions influencing animal movement, mating structure, and selection intensity, following approaches used in prior native chicken and smallholder livestock studies [10,11,12].

### Phenotypic measurements

Phenotypic variation was assessed using both morphometric and plumage-colour traits measured on individual chickens. Morphometric measurements comprised body height (vertical distance from the ground to the dorsal body line), wingspan (distance between the tips of the fully extended wings), chest width (widest region of the sternum), shank length (distance along the tarsometatarsus between the proximal and distal articulations), and shank diameter (cross-sectional dimension of the tarsometatarsus at its midpoint, recorded along the mediolateral [short axis] and anteroposterior [long axis] planes). All measurements (cm) were obtained using a measuring tape or digital caliper, following standardized protocols with calibrated instruments to minimize inconsistencies across sampling locations.

Plumage colour was quantified from RGB values recorded for the head, body, and tail regions of each chicken. Because RGB values are device-oriented and highly correlated, they were converted to hue–saturation–value (HSV) colour space prior to analysis, with hue represented using circular coordinates (cosine and sine transformations) to avoid discontinuities associated with its circular nature. The final dataset therefore included hue coordinates, saturation, and value for each plumage region.

### Data structure and spatial grouping

Data were organized hierarchically, with individual chickens nested within sampling locations, locations within island systems, and island systems within provinces — reflecting the geographic configuration of the Wallacea archipelago and enabling phenotypic variation to be evaluated across local, island, and regional spatial scales.

Because island and province were not fully nested, island-level variance differed in interpretation across regions: within Wallacea, islands provided resolution below the province level (multiple islands per province), whereas the three Java provinces shared a single island grouping, so island-level variance for Java reflects variation shared across provinces rather than sub-province structure. Hierarchical mixed-effects models accommodate this type of unbalanced multilevel data without requiring a fully balanced design [20,21].

### Multivariate analysis of phenotypic variation

Principal component analysis (PCA) was used to summarize multivariate phenotypic variation and reduce dimensionality, run separately for (i) morphometric and (ii) plumage-colour traits after standardizing all variables to zero mean and unit variance [22,23].

For each PCA, the variance explained by each component was recorded, and the first two components were used to visualize phenotypic patterns across provinces and island systems, assessing multivariate overlap, dispersion, and spatial structuring. For plumage colour, PCA was conducted on HSV-derived variables for head, body, and tail plumage only; shank and comb colour were excluded to focus specifically on feather colour variation.

### Quantification of phenotypic structure

To quantify the relative contribution of hierarchical spatial scales to phenotypic variation, variance components were estimated separately for each of the six morphometric traits (body height, wingspan, chest width, shank length, and shank diameter at the short and long axes) using linear mixed-effects models (lme4 package), with island, province, and sampling location nested within province included as random effects. The model for each trait was specified as *Y* = *μ* + *Island* + *Province* + *Location*(*Province*) + *e*, where Y represents the individual morphometric measurement, μ is the overall mean, and e represents residual variation; models were fitted using restricted maximum likelihood (REML). Variance components were extracted and expressed as percentages of total variance, partitioning it into components attributable to islands, provinces, locations within provinces, and residual (within-location) variation [20,21]. PCA scores from morphometric traits were analyzed using the same hierarchical structure to evaluate whether multivariate patterns were consistent with variance partitioning from individual traits.

Because East Nusa Tenggara showed unusually high dispersion in selected morphometric traits, sensitivity analyses excluding this province were conducted — repeating PCA, variance partitioning, and PERMANOVA on the reduced dataset — to evaluate whether the overall interpretation was robust to potential location-specific effects.

### Multivariate analysis of spatial structure

Multivariate phenotypic structure was further assessed using permutational multivariate analysis of variance (PERMANOVA [24]) on Euclidean distance matrices calculated from standardized morphometric traits, evaluating the contribution of island, province, and sampling location to morphometric differentiation (999 permutations, sequential term tests). An additional model with island alone evaluated the overall island effect. PERMANOVA provides a flexible framework for assessing the extent to which geographic and hierarchical spatial factors contribute to overall phenotypic variation in observational datasets.

### Analytical framework

Given the observational nature of the data and the absence of predefined breeding populations, the analytical approach quantified phenotypic structure rather than inferring population genetic parameters directly, guided by frameworks relating variation to the expected effects of gene flow, drift, and selection [3,4,25]. Genotype-by-environment interaction was not treated as a discriminant factor here. All statistical analyses were performed in R [26], using the lme4 and vegan packages for mixed-effects modelling and PERMANOVA, respectively.

## Results

The final dataset comprised 612 chickens sampled from 259 households across eight provinces spanning Java and the Wallacea region, encompassing multiple island systems and a broad range of agroecological conditions.

### Production system characteristics

Production systems differed between Java and Wallacea while retaining key features of smallholder chicken production (Table 3). Flock sizes were generally small across all regions (approximately 12–24 birds per household), with chickens predominantly managed under free-range or semi-scavenging systems, particularly in Wallacea, where birds commonly roamed freely around households.

**Table 3.** Summary of production system characteristics across regions.

| Variable | Wallacea (range across locations) | Java (range across locations) |
| --- | --- | --- |
| Flock size (birds/household, mean ± SD) | 12.8–23.9 (± 8.8–23.5) | 14.6–18.8 (± 7.0–14.5) |
| Rearing system (% free-range without fencing) | Predominantly high (43.5–92.5%) | Lower to moderate (0–70%) |
| Housing availability (% with shelter) | Limited (0–50%) | More common (16.7–75%) |
| Housing structure | Mostly open (no roof/wall dominant) | More structured (roof and wall present) |
| Feeding practice (% supplementary feeding) | Universal, but largely non-commercial (grain, waste, mixed feed) | Universal, with higher use of commercial feed |
| Access to forage (%) | High (72–100%) | High (91.7–100%) |
| Animal exchange / replacement | Frequent local exchange and informal transfer | Local exchange + greater market involvement |
| Marketing channels | Predominantly direct sale and traditional markets | More diversified (direct, market, middleman) |

Values are aggregated from household survey data across locations within Wallacea and Java.
Ranges indicate variation among provinces within each region.

Housing and feeding practices varied among locations: more structured housing and greater commercial feed use were observed in Java, while Wallacea relied more on scavenging and locally available feed. Flock replacement and animal exchange occurred primarily through informal mechanisms, local markets, neighbour-to-neighbour transfer, cultural functions, direct sale, and within-flock replacement, indicating continued movement of chickens across communities despite geographic fragmentation among island systems.

### Farmer-defined objectives and selection practices

Farmers reported multiple and overlapping objectives for keeping chickens, including household consumption, income generation, and socio-cultural purposes (Table 4). No single production objective consistently dominated across regions, and motivations for maintaining chickens varied among households and locations.

**Table 4.** Farmer-defined production objectives, motivations, and selection criteria across regions.

| Category | Trait / Objective | Wallacea (range) | Java (range) |
| --- | --- | --- | --- |
| Production function | Meat production (household consumption) | High (0.315–0.405) | Moderate (0.208–0.400) |
|  | Egg production (household consumption) | Moderate (0.136–0.310) | Moderate to high (0.250–0.375) |
|  | Income generation | Moderate to high (0.230–0.373) | Moderate to high (0.260–0.431) |
|  | Socio-cultural / ritual | Low to moderate (0.057–0.176) | Low (0.017–0.156) |
| Motivation to keep chickens | Food waste utilization | High (0.164–0.401) | Moderate to high (0.153–0.404) |
|  | Income generation | Moderate (0.219–0.333) | Moderate (0.278–0.404) |
|  | Pet / non-productive use | Moderate to high (0.234–0.433) | Variable (0.105–0.431) |
| | Hobby / competition | Low ($\leq 0.199$) | Low ($\leq 0.088$) |
| Male selection criteria | Body size / weight | Dominant (0.440–0.473) | Dominant (0.444–0.469) |
|  | Plumage color | Moderate importance (0.258–0.367) | Moderate importance (0.222–0.317) |
|  | Comb morphology | Secondary (0.084–0.187) | Secondary to moderate (0.100–0.240) |
| Female selection criteria | Body size / weight | Dominant (0.387–0.457) | Dominant (0.375–0.472) |
|  | Egg production | Moderate to high (0.269–0.333) | High (0.194–0.427) |
|  | Plumage color | Secondary (0.110–0.213) | Secondary (0.062–0.233) |
|  | Other traits | Minor importance | Minor importance |

Values represent rank index scores derived from household surveys. Ranges indicate variation across locations within Wallacea and Java.

Selection of breeding animals was primarily based on observable phenotypic characteristics. For males, body size and general appearance were consistently prioritized, followed by plumage and comb morphology; for females, body size and egg production were the most frequently reported criteria. Relative importance varied among regions and households, reflecting heterogeneous farmer preferences and decentralized management objectives.

### Morphometric variation across locations

Substantial variation was observed across all measured morphometric traits (Table 5). Mean values differed among provinces and island systems for body size and extremity measurements, including body height, wingspan, chest width, and shank dimensions. East Nusa Tenggara showed particularly high mean values for wingspan and shank length, whereas differences among other regions were generally more moderate.

**Table 5.** Morphometric variation of local chickens across sampling locations (mean ± SD).

| Province | Island system | n | Body height<br>(cm) | Wingspan<br>(cm) | Chest width<br>(cm) | Shank length<br>(cm) | Shank diameter<br>(short, cm) | Shank diameter<br>(long, cm) |
| --- | --- | --- | --- | --- | --- | --- | --- | --- |
| Central Java | Java | 40 | 28.68 ± 2.30 | 31.77 ± 2.63 | 7.24 ± 0.83 | 6.07 ± 0.83 | 0.96 ± 0.10 | 1.27 ± 0.13 |
| East Java | Java | 36 | 29.28 ± 1.98 | 32.14 ± 2.85 | 7.13 ± 0.95 | 6.01 ± 0.45 | 0.96 ± 0.13 | 1.22 ± 0.15 |
| West Java | Java | 63 | 26.71 ± 1.78 | 31.27 ± 1.89 | 6.31 ± 0.77 | 5.67 ± 0.52 | 0.86 ± 0.07 | 1.11 ± 0.13 |
| Maluku | Ambon–Seram islands | 100 | 27.90 ± 2.80 | 31.72 ± 2.72 | 6.57 ± 0.88 | 5.72 ± 0.76 | 0.85 ± 0.15 | 1.22 ± 0.19 |
| North Maluku | Halmahera–Ternate<br>islands | 100 | 27.11 ± 1.93 | 33.45 ± 2.75 | 6.73 ± 0.98 | 6.10 ± 0.74 | 0.88 ± 0.08 | 1.17 ± 0.11 |
| Southeast<br>Sulawesi | Buton–Muna islands | 74 | 26.70 ± 3.20 | 35.01 ± 3.24 | 7.37 ± 0.90 | 5.82 ± 0.64 | 0.94 ± 0.11 | 1.27 ± 0.15 |
| West Nusa<br>Tenggara | Lombok–Sumbawa<br>islands | 107 | 26.58 ± 2.34 | 32.13 ± 2.02 | 6.12 ± 0.70 | 6.79 ± 0.57 | 0.92 ± 0.11 | 1.25 ± 0.13 |
| East Nusa<br>Tenggara | Timor–Flores–Sumba<br>islands | 92 | 24.46 ± 2.61 | 41.23 ± 3.49 | 6.43 ± 1.55 | 8.43 ± 0.77 | 1.04 ± 0.83* | 1.17 ± 0.26 |

Despite these differences, variation within regions remained considerable, and ranges of trait values overlapped extensively among provinces and island systems, with individuals from different geographic and island contexts frequently sharing similar morphometric characteristics, indicating that phenotypic variation was distributed continuously rather than forming discrete regional types. Variation among traits was also heterogeneous: province-level differences appeared more pronounced for wingspan and shank length, whereas other traits, particularly shank diameter, showed substantial within-region variation relative to differences among geographic groups.

### Multivariate phenotypic structure

Principal component analysis of morphometric traits revealed limited separation among populations in multivariate space (Fig. 2). Individuals from different provinces and island systems occupied broadly overlapping regions, with no clear clustering corresponding to geographic origin. Chickens from East Nusa Tenggara tended to occupy a more dispersed and partially separated region of the PCA space, though substantial overlap with other provinces remained.

**Fig 1.**
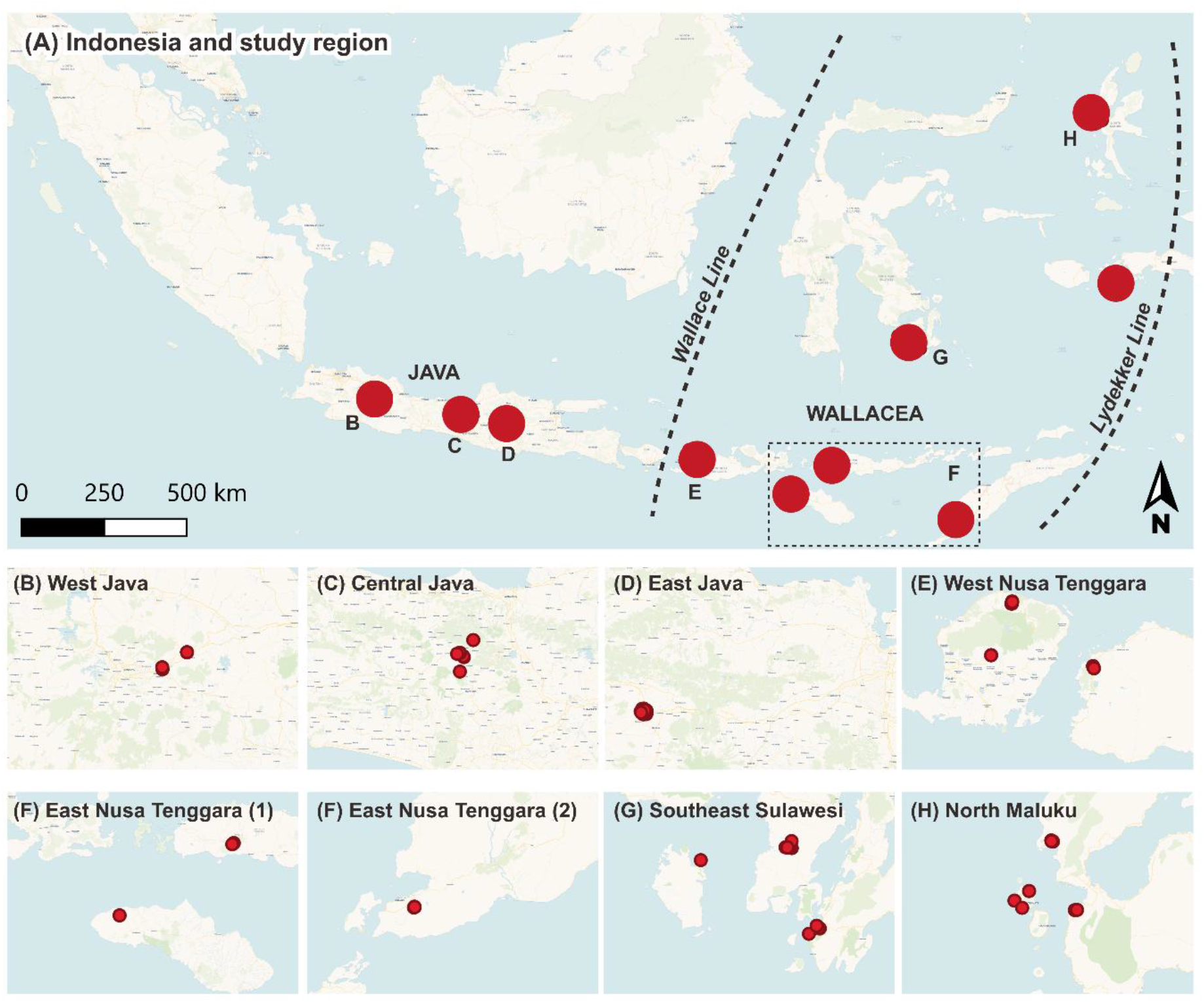
Sampling locations across Indonesia and the Wallacea region. (A) Geographic distribution of sampling sites across Java and Wallacea, showing major biogeographic boundaries (Wallace and Lydekker Lines); red points indicate sampling locations, dashed boxes highlight sampled regions. (B–H) Detailed views within each province: West Java (B), Central Java (C), East Java (D), West Nusa Tenggara (E), East Nusa Tenggara (F), Southeast Sulawesi (G), North Maluku (H).

**Fig 2.**
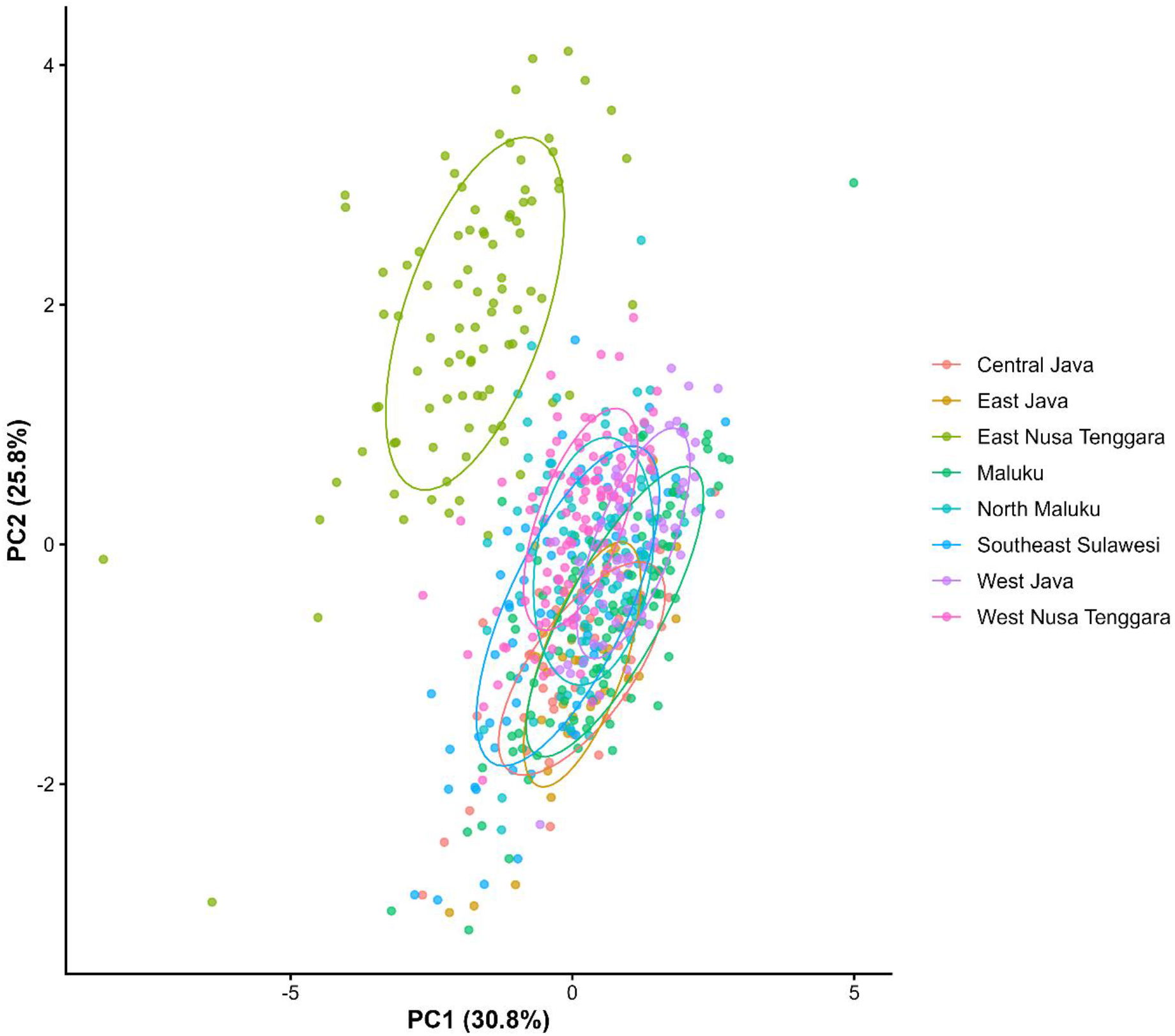
Multivariate structure of morphometric traits across sampling locations, standardized using PCA. Points represent individual chickens, coloured by province; ellipses indicate within-province dispersion. PC1 and PC2 are shown, with variance explained indicated on each axis.

The first two principal components captured broad gradients of morphometric variation rather than discrete regional structure, so differences among provinces and island systems were expressed mainly as shifts in the distribution and dispersion of individuals rather than sharply differentiated clusters. These results were consistent with the descriptive morphometric analyses (Table 5), together indicating that morphometric diversity was distributed continuously across populations despite geographic fragmentation across the Wallacea archipelago.

### Quantification of phenotypic structure

Quantitative analyses supported the multivariate patterns. Variance partitioning showed that most phenotypic variation occurred within locations, though the relative contribution of higher spatial levels differed among traits (Table 6; Fig. 3), within-location variation remained the dominant component across traits, indicating substantial diversity maintained locally despite geographic and island fragmentation.

**Fig 3.**
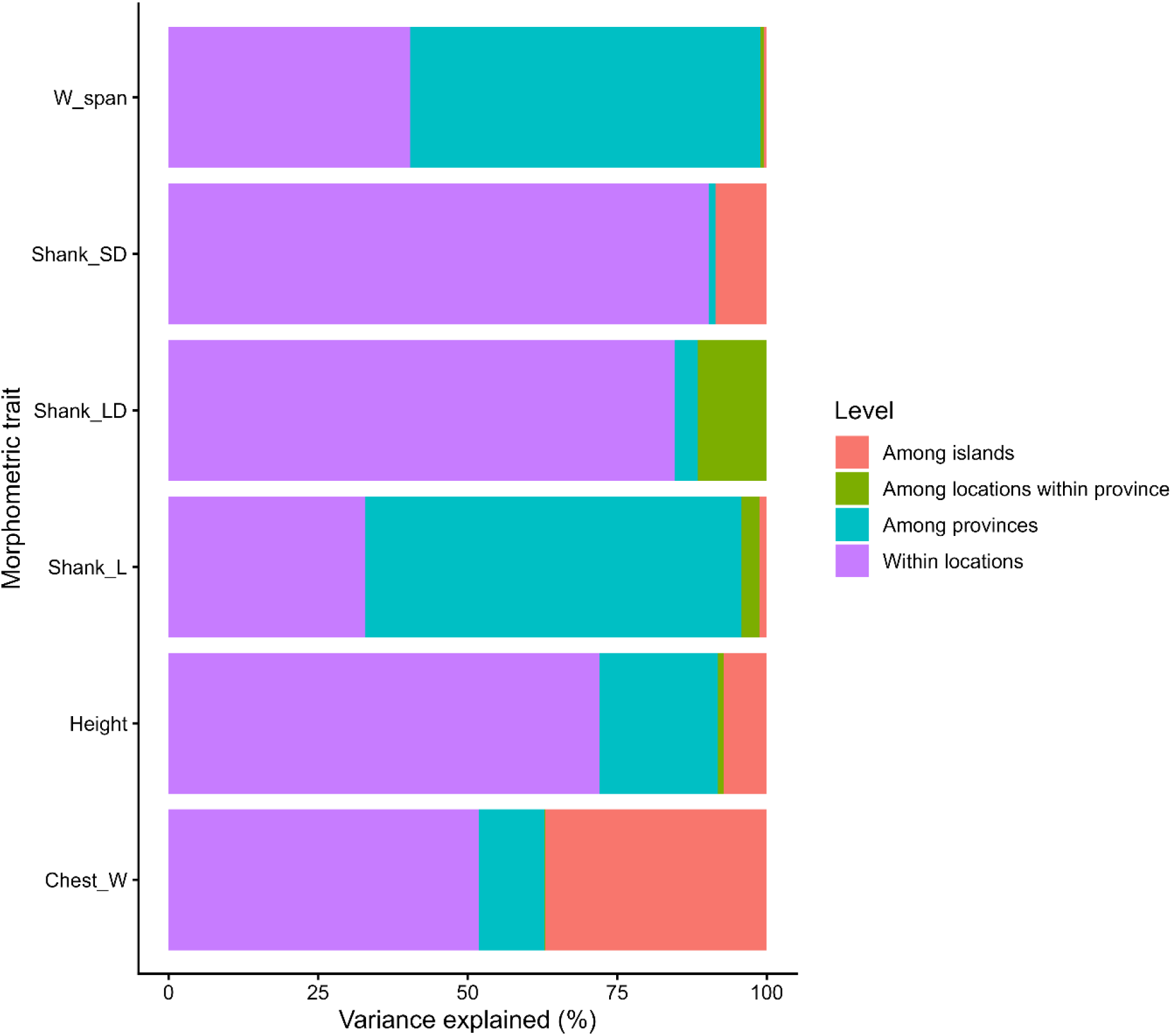
Variance partitioning of morphometric traits across spatial scales: proportion of total phenotypic variance attributable to within-location variation, among-location (within-province), and among-province differences, estimated from linear mixed-effects models and expressed as percentages.

**Table 6.** Variance partitioning of morphometric traits across hierarchical spatial scales including island structure.

| Trait | Within locations (%) | Among provinces (%) | Among islands (%) | Among locations within province (%) |
| --- | --- | --- | --- | --- |
| Body height | 72.02 | 19.83 | 7.17 | 0.98 |
| Wingspan* | 40.39 | 58.56 | 0.41 | 0.63 |
| Chest width* | 51.91 | 10.93 | 37.07 | 0.10 |
| Shank length* | 32.84 | 62.94 | 1.19 | 3.03 |
| Shank diameter (short) | 90.35 | 1.08 | 8.57 | <0.01 |
| Shank diameter (long) | 84.62 | 3.89 | <0.01 | 11.49 |

Values represent the proportion (%) of total phenotypic variance attributable to within-location variation, differences among provinces, differences among islands, and differences among locations within provinces for each morphometric trait.

The contribution of island-and province-level structure varied among traits: province-level effects were most pronounced for wingspan and shank length, while island-level effects contributed appreciably to chest width and, to a lesser extent, shank short diameter (Table 6; sensitivity to East Nusa Tenggara examined below and in Supplementary Table S1–S2). Differences among locations within provinces generally accounted for smaller proportions of total variance. PERMANOVA further indicated that spatial grouping explained a significant but moderate proportion of total morphometric variation (R² = 0.361, P < 0.001); despite this detectable geographic structure, individuals from different provinces and island systems continued to show extensive multivariate overlap (Fig. 2), indicating diffuse, continuously distributed rather than discretely partitioned phenotypic variation.

Sensitivity analyses excluding East Nusa Tenggara showed that the overall pattern of diffuse, within-location-dominated phenotypic structure was preserved, and for several traits even more pronounced, though the magnitude of province-and island-level effects for specific traits was not robust to this exclusion (Supplementary Fig. S1–S3; Supplementary Table S1–S2). Within-location variation increased for most traits once East Nusa Tenggara was excluded (wingspan: 40.4% to 80.0%; shank length: 32.8% to 71.3%; chest width: 51.9% to 73.1%), while the province-level effects previously identified for wingspan and shank length, and the island-level effect for chest width, were substantially reduced (wingspan among-province variance: 58.6% to 15.2%; shank length: 62.9% to 22.9%; chest width among-island variance: 37.1% to 4.7%). East Nusa Tenggara therefore contributed disproportionately to the higher-level spatial structuring detected for these three traits, while the broader pattern of extensive within-location variation and diffuse overlap among provinces and islands (Fig. 2) did not depend on this single location.

### Plumage color variation

PCA of HSV-derived plumage colour traits revealed continuous but diffuse variation across populations (Fig. 4): PC1 explained 57.2% of total variation and PC2 explained 15.7%, indicating that plumage colour variation was structured primarily along a dominant but non-exclusive axis of differentiation.

**Fig 4.**
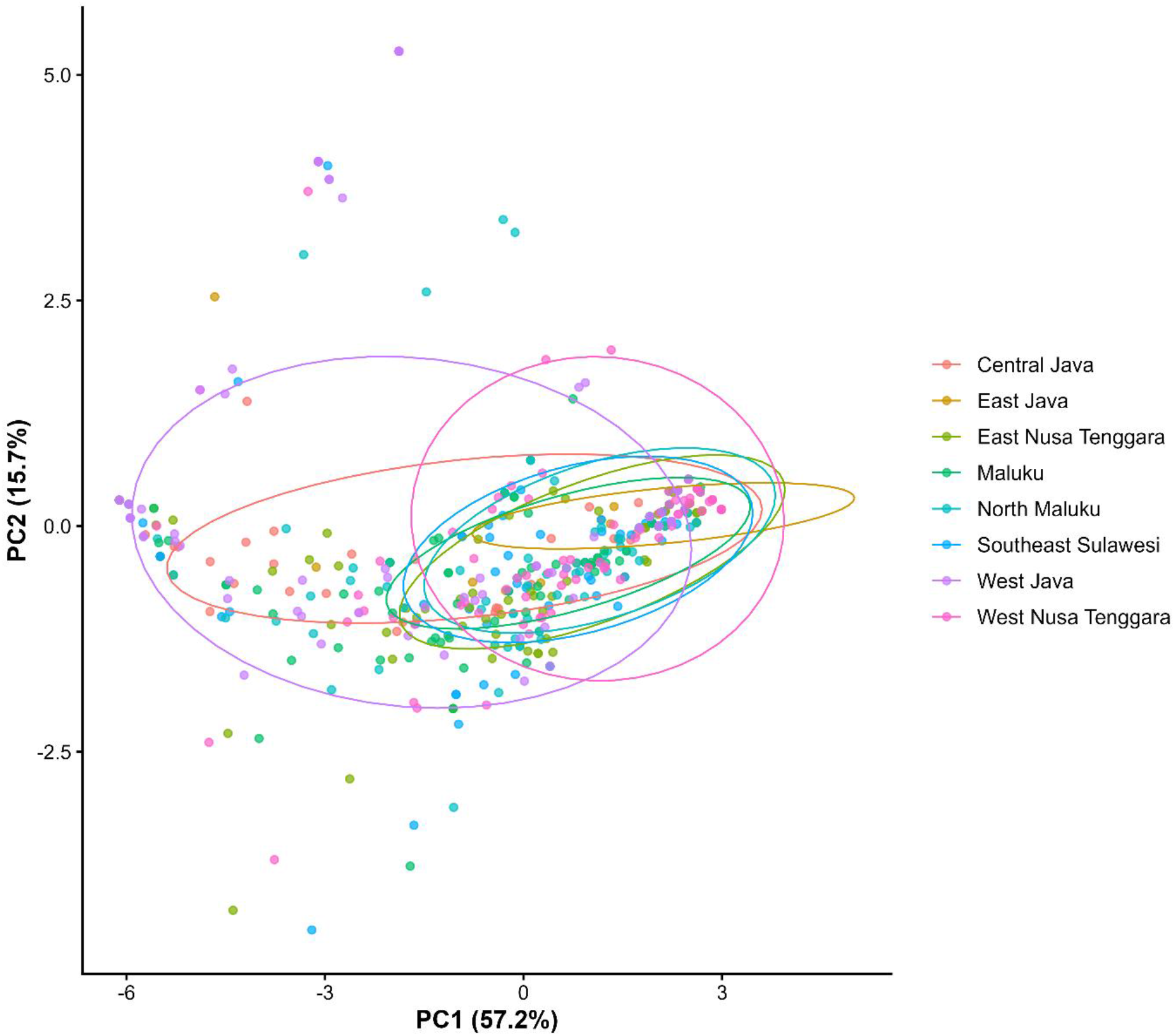
Multivariate structure of plumage colour variation based on HSV-derived traits for head, body, and tail plumage (hue represented using circular coordinates). Points represent individual chickens coloured by province; ellipses indicate within-province dispersion. PC1 and PC2 explained 57.2% and 15.7% of total variation, respectively.

Individuals from different provinces and island systems showed substantial overlap in plumage-colour space, with no clear separation into discrete regional clusters. Some provinces displayed broader dispersion along PC1, indicating differences in the frequency and range of plumage phenotypes rather than the presence of province-specific colour types. As with the morphometric analyses, plumage phenotypes were characterised more by overlapping gradients of variation than by sharply differentiated regional groupings.

Across both morphometric and plumage-colour traits, phenotypic variation was characterized by extensive within-location diversity combined with weak and diffuse spatial structuring. Although province and island-level effects were detectable for selected morphometric traits (Table 6; Fig. 3), these differences did not produce discrete phenotypic clusters in multivariate space (Figs. 2 and 4).

Individuals from different provinces and island systems consistently showed broad overlap in both morphometric and plumage-colour variation, so phenotypic structure appeared continuous and hierarchical rather than sharply partitioned into distinct regional types, indicating that native chicken populations across Wallacea and Java retain substantial local variation despite geographic fragmentation.

## Discussion

The phenotypic patterns observed here must be considered alongside the production systems in which these chickens are kept. Extensive overlap among locations, the predominance of within-location variation, continued exchange of chickens, and heterogeneous farmer selection together indicate a decentralized production system with substantial connectivity but limited coordination of selection — human-mediated exchange offering a plausible mechanism through which animals, and potentially genes, move among locations. These remain mechanistic hypotheses rather than direct estimates of gene flow.

### Phenotypic variation under diffuse spatial structuring

The results reveal a consistent pattern of high within-location variation combined with weak but detectable spatial structuring. Morphometric traits showed extensive overlap among provinces in multivariate space (Fig. 2), and variance partitioning indicated that most phenotypic variation was maintained within locations rather than among them (Table 6), indicating hierarchical but diffuse rather than sharply partitioned structuring.

This pattern contrasts with classical population genetic theory, where geographic separation and limited dispersal are predicted to generate stronger differentiation [3,4]. Instead, the observed structure reflects a balance between processes promoting differentiation and those maintaining connectivity: gene flow can limit local divergence when migration counteracts spatially divergent selection [27,28], so the weak structuring here is consistent with substantial human-mediated connectivity, though the data do not directly estimate gene flow or selection strength. Sensitivity analyses excluding East Nusa Tenggara reinforced this, though province-and island-level effects for three traits were substantially reduced, indicating this location contributed disproportionately to higher-level structuring in the full dataset.

High levels of within-location variation are not merely residual noise but a fundamental component of evolutionary systems: contemporary theory emphasizes that substantial phenotypic diversity can be maintained within populations under gene flow, drift, and heterogeneous selection [29], reflecting genuine local diversity rather than an absence of spatial structure.

### Production systems as drivers of connectivity and potential gene flow

The weak spatial structuring aligns with connectivity patterns typical of smallholder chicken systems. Survey data indicated frequent chicken exchange among households through informal transfer, local markets, and within-community replacement (Table 2); combined with free-range management and uncontrolled mating, these practices likely facilitate animal movement and gene flow, though this is not directly estimated here.

Connectivity is not determined solely by geographic proximity: market access, transport routes, topography, and inter-island trading can facilitate or constrain exchange, so nearby groups may become partially isolated while distant groups remain connected, weakening expected effects of geographic separation in Wallacea.

These dynamics differ from natural populations, where dispersal is more constrained by ecological and geographic barriers [5,6]. In domesticated populations, continued human-mediated exchange can maintain connectivity and facilitate gene flow, consistent with the extensive overlap observed here and the modest island-level effects for most traits.

Similar patterns occur in village chicken systems across Africa and Asia, where high within-population variation and limited among-population differentiation are commonly observed under decentralized management [13,14,15], supporting connectivity and animal exchange as system-level drivers of phenotypic diversity; whether this reflects contemporary gene flow requires genomic evaluation.

### Decentralized and weakly coordinated selection under smallholder conditions

A second key feature of the system is farmer-driven selection. Selection criteria were primarily based on observable traits, body size and plumage characteristics, but applied heterogeneously and decentrally across households (Table 3), with farmers reporting multiple and sometimes competing objectives (meat and egg production, income generation, non-productive uses), resulting in a diffuse and inconsistent selection regime.

Under such conditions, selection is less likely to generate consistent directional change, since preferences vary among households; heterogeneous selection may instead help maintain diverse phenotypes while exchange and demographic processes reduce differentiation. The HSV-based plumage-colour analysis supports this: colour varied along major axes, yet individuals from different provinces overlapped extensively, visible traits differ mainly in frequency and dispersion, varying continuously rather than forming discrete regional phenotypes.

This suggests that selection on observable phenotypes operates at a local scale but is insufficiently coordinated to produce stable regional differentiation, its effects likely counteracted by ongoing animal exchange and uncontrolled mating. Similar conclusions have been drawn in smallholder livestock systems, where decentralized decision-making and limited information constrain the effectiveness of selection [10,12].

### Interaction between connectivity, drift, and decentralized selection

The observed patterns likely reflect interacting evolutionary, environmental, and management processes: animal exchange may maintain connectivity among households, small flock sizes may increase the influence of genetic drift, and decentralized farmer selection varies in its preferences for body size, plumage, and production traits, together explaining why substantial variation is maintained within locations while broader differentiation remains limited.

These processes operate across agroecologically diverse, fragmented island systems. Although island-level effects were detectable for chest width, structure remained diffuse overall, indicating environmental heterogeneity and fragmentation alone are insufficient to generate discrete structuring under ongoing connectivity. Geographic separation does not necessarily mean biological isolation: human-mediated exchange through trading, markets, and inter-island movement may maintain connectivity even as topography and market access shape animal movement.

These findings are consistent with theoretical expectations that migration, drift, and selection operate simultaneously across heterogeneous environments [28], interpreted here as potential mechanisms rather than directly estimated parameters, genomic data will be required to confirm whether contemporary gene flow contributes to the pattern observed.

### Implications for biogeography and domesticated populations

The lack of strong phenotypic structuring highlights the limited, trait-dependent role of biogeographic barriers in shaping domesticated populations under smallholder management.

Wallacea is well known for pronounced genetic differentiation in wild species [16], driven by historical isolation and restricted dispersal [30], yet this pattern is not necessarily reflected in domesticated chickens, which may be strongly influenced by human-mediated movement, so patterns may reflect contemporary system dynamics rather than historical biogeographic boundaries [18,19].

### Broader implications for animal genetic resources

More broadly, animal genetic resources in smallholder systems are best understood as dynamic and interconnected rather than discrete, geographically bounded populations. Animal exchange, small flock sizes, and weakly coordinated selection together may shape the maintenance and redistribution of phenotypic diversity, so breeding and conservation approaches assuming well-defined populations may be poorly suited here; effective interventions should account for farmer practices, exchange networks, and decentralized selection, with relevance extending to smallholder livestock systems generally in low-and middle-income countries.

### Limitations and future directions

Several limitations should be considered. First, the study relied primarily on phenotypic and household-level data and cannot directly distinguish among the evolutionary mechanisms underlying the observed patterns. The diffuse structure is consistent with human-mediated connectivity, potential gene flow, and weakly coordinated selection, but similar patterns could also arise through retained ancestral variation or convergent environmental responses; the overlap observed should therefore be read as evidence of diffuse differentiation, not direct evidence of gene flow.

Second, East Nusa Tenggara showed unusually high dispersion for selected traits. Sensitivity analyses confirmed the central conclusion of diffuse, within-location-dominated variation was robust, and strengthened, once this province was excluded, though province-and island-level effects for wingspan, shank length, and chest width were substantially attenuated (see Results). Whether this reflects genuine biological distinctiveness, greater local heterogeneity, or measurement artifact (Table 5) warrants targeted follow-up sampling.

Third, analyses were restricted to adult female chickens to reduce confounding from sex-specific ornamental traits, trading practices, and non-random male exchange. Male chickens may experience different selection pressures related to body size, plumage, reproductive roles, or cultural preferences; including both sexes in future studies would clarify sex-specific contributions to phenotypic structuring.

Finally, genomic data will be required to evaluate the genetic mechanisms underlying these patterns. Ongoing genomic analyses of these populations will help distinguish among historical ancestry, contemporary gene flow, and local adaptation, and assess whether genetic connectivity corresponds to the animal-exchange networks identified here, clarifying the relative contributions of connectivity and selection to chicken diversity across Wallacea and Java.

## Conclusion

Native chickens across Wallacea and Java exhibited substantial phenotypic diversity characterized by extensive within-location variation and diffuse spatial structuring. Although province-and island-level effects were detectable for selected morphometric traits, individuals from different provinces and islands showed broad overlap in morphometric and plumage-colour variation, indicating continuous rather than discrete differentiation. These patterns are consistent with decentralized smallholder production, heterogeneous management, weakly coordinated selection, and continued human-mediated exchange, suggesting geographic fragmentation and agroecological heterogeneity alone are insufficient to generate strongly differentiated phenotypic groups. Whether this connectivity reflects contemporary gene flow, retained ancestral variation, or convergent environmental responses remains to be determined through genomic analysis.

## Ethics approval

All procedures involving animals were approved by the Animal Ethics Committee, Faculty of Animal Science, Universitas Sebelas Maret (Approval No. 150.2/UN27.14/TA.00.03/2026, dated 6 February 2026), and were conducted in accordance with relevant institutional and national guidelines for the care and use of animals. The study was observational and involved non-invasive measurements of animals under field conditions.

## Consent to participate

Informed consent was obtained from all participating farmers prior to data collection.

## Data and model availability statement

The datasets generated and analyzed during the current study are available from the corresponding author upon reasonable request.

## Declaration of generative AI and AI-assisted technologies in the writing process

During the preparation of this work, the authors used Scopus AI for reference searches and Claude to enhance grammar, and language clarity. After using these tools, the authors reviewed and edited the content as needed and take full responsibility for the content of the published article.

## Supporting information

https://drive.google.com/drive/folders/1QZWeBSXEfQmPtkHpWhTu6Z1vNtG8egF8?usp=sharing

## Acknowledgements

The authors sincerely thank all farmers and local communities involved in this study for their cooperation during field sampling and interviews across Java and Wallacea, and acknowledge the assistance of field enumerators, students, and collaborating institutions who contributed to data collection and logistical support. The authors are especially grateful to Professor Olivier Hanotte for his valuable comments and constructive suggestions during the revision process.

## Author approval and publication status

All authors have seen and approved the final version of this manuscript and agree with its submission to bioRxiv. This manuscript has not been accepted for publication and has not been published elsewhere.

