## Supplementary material for "Diffuse phenotypic structure in native chickens across Java and Wallacea is consistent with human-mediated connectivity and decentralized selection": https://drive.google.com/drive/folders/1QZWeBSXEfQmPtkHpWhTu6Z1vNtG8egF8?usp=sharing

**Supplementary Materials**

**Sensitivity analysis excluding East Nusa Tenggara (island-informed)**

East Nusa Tenggara showed unusually high dispersion in several morphometric traits, including wingspan and shank measurements (Table 5). To evaluate whether the overall interpretation of phenotypic structure was robust to this province, sensitivity analyses were repeated after excluding all East Nusa Tenggara chickens (n = 92 of 612), reducing the dataset to 520 individuals across seven provinces. Principal component analysis (PCA), variance partitioning using linear mixed-effects models, and permutational multivariate analysis of variance (PERMANOVA) were repeated on this reduced dataset using the same island-informed model structure (Island, Province, and Location within Province) used for the full dataset (Table 6).

The sensitivity analyses confirmed that the central conclusion of diffuse, within-location-dominated phenotypic variation was robust to the exclusion of East Nusa Tenggara, and in several cases became more pronounced. Mean within-location variance across the six morphometric traits increased from 62.0% in the full dataset to 78.6% after exclusion of East Nusa Tenggara, while mean among-province variance decreased from 26.2% to 15.8% and mean among-island variance decreased from 9.1% to 2.4% (Supplementary Table S1; Supplementary Fig. S3). No discrete phenotypic clustering emerged in PCA space after exclusion of East Nusa Tenggara (Supplementary Fig. S1), consistent with the pattern observed in the full dataset (Fig. 2).

However, this robustness was not uniform across traits. The province-level effects previously identified as pronounced for wingspan and shank length were substantially reduced after excluding East Nusa Tenggara (wingspan: 58.6% to 15.2%; shank length: 62.9% to 22.9%), and the island-level effect identified for chest width was similarly reduced (37.1% to 4.7%); within-location variance increased correspondingly for these three traits (wingspan: 40.4% to 80.0%; shank length: 32.8% to 71.3%; chest width: 51.9% to 73.1%). This indicates that East Nusa Tenggara contributed disproportionately to the higher-level spatial structuring detected for these three traits in the full-dataset analysis. In contrast, shank diameter (short and long) and body height showed comparatively modest changes upon exclusion of this province (Supplementary Table S1; Supplementary Fig. S2).

PERMANOVA results were consistent with this pattern. Spatial grouping explained a smaller proportion of total morphometric variation after exclusion of East Nusa Tenggara (Island + Province + Location: $R^2 = 0.210$, compared with $R^2 = 0.361$ in the full dataset), although Island,

Province, and Location each remained statistically significant contributors to multivariate phenotypic structure ($P \leq 0.002$ for all terms; Supplementary Table S2).

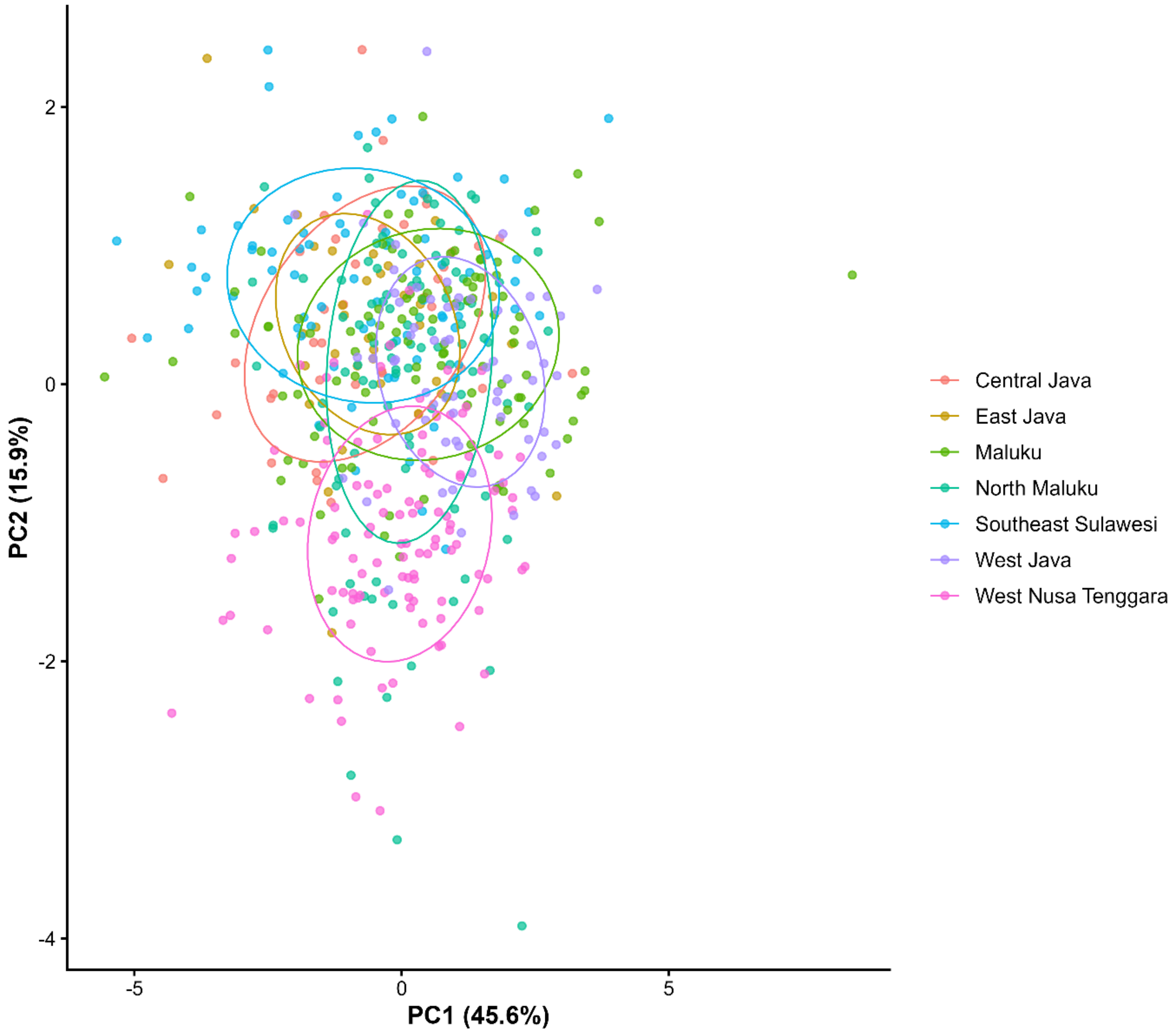


**Supplementary Figure S1. Sensitivity analysis of morphometric PCA excluding East Nusa Tenggara.**

Principal component analysis (PCA) of standardized morphometric traits after exclusion of East Nusa Tenggara. Points represent individual chickens coloured by province; ellipses indicate the dispersion of individuals within each province. No discrete phenotypic clustering emerged, consistent with the diffuse structuring observed in the full dataset.

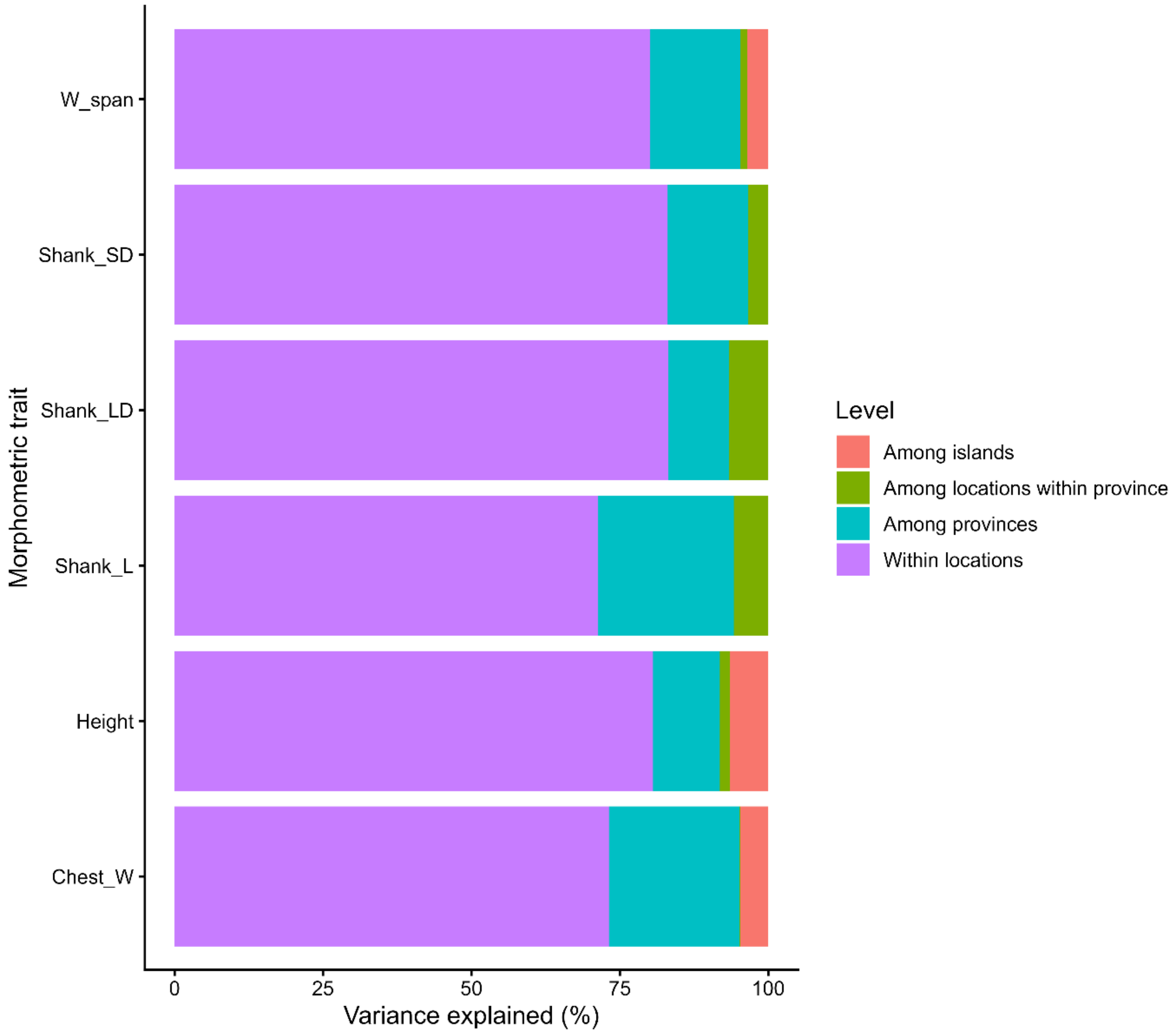


**Supplementary Figure S2. Variance partitioning of morphometric traits excluding East Nusa Tenggara.**

Proportion of total morphometric variance attributable to within-location variation, among-province variation, among-island variation, and among-location-within-province variation after exclusion of East Nusa Tenggara, estimated using the same island-informed linear mixed-effects models used for the full dataset (Table 6). Within-location variation remained the dominant component for most traits and increased for wingspan, shank length, and chest width relative to the full dataset (Supplementary Table S1).

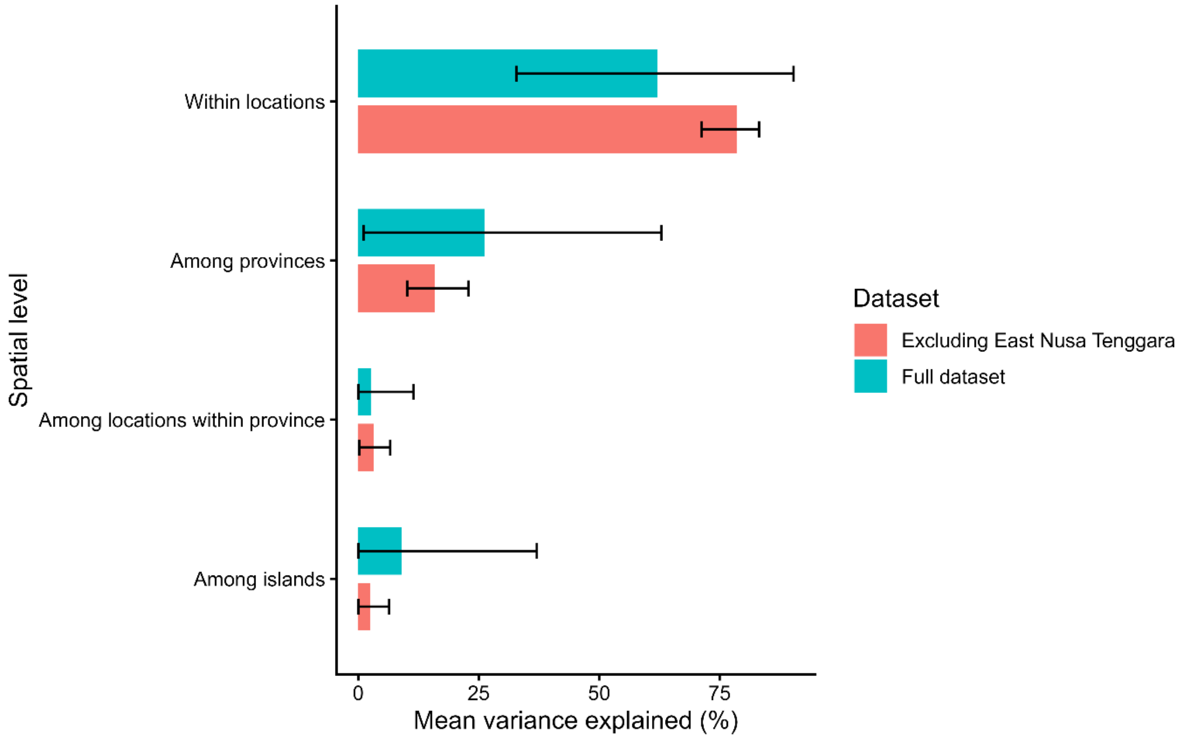


**Supplementary Figure S3. Comparison of variance partitioning between the full dataset and the dataset excluding East Nusa Tenggara.** Mean proportion of morphometric variance (± range across the six traits) attributable to each spatial level in the full dataset and in the sensitivity analysis excluding East Nusa Tenggara. Within-location variance increased and among-province and among-island variance decreased on average after exclusion of this province; trait-specific results are given in Supplementary Table S1.

**Supplementary Tables**

**Supplementary Table S1. Variance partitioning of morphometric traits excluding East Nusa Tenggara, compared with the full dataset.**

| Trait | Spatial level | Full dataset (%) | Excl. E. Nusa Tenggara (%) | Δ (pp) |
|---|---|---|---|---|
| Body height | Within locations | 72.02 | 80.60 | +8.58 |
| | Among provinces | 19.83 | 11.23 | -8.60 |
| | Among islands | 7.17 | 6.42 | -0.75 |
| | Among locations within province | 0.98 | 1.75 | +0.77 |
| Wingspan | Within locations | 40.39 | 80.03 | +39.64 |
| | Among provinces | 58.56 | 15.22 | -43.34 |
| | Among islands | 0.41 | 3.54 | +3.13 |
| | Among locations within province | 0.63 | 1.21 | +0.58 |
| Chest width | Within locations | 51.91 | 73.15 | +21.24 |
| | Among provinces | 10.93 | 21.98 | +11.05 |
| | Among islands | 37.07 | 4.66 | -32.41 |
| | Among locations within province | 0.10 | 0.22 | +0.12 |
| Shank length | Within locations | 32.84 | 71.29 | +38.45 |
| | Among provinces | 62.94 | 22.90 | -40.04 |
| | Among islands | 1.19 | 0.00 | -1.19 |
| | Among locations within province | 3.03 | 5.81 | +2.78 |
| Shank diameter (short) | Within locations | 90.35 | 83.07 | -7.28 |
| | Among provinces | 1.08 | 13.56 | +12.48 |
| | Among islands | 8.57 | 0.00 | -8.57 |
| | Among locations within province | 0.00 | 3.37 | +3.37 |

| | | | | |
|---|---|---|---|---|
| Shank diameter (long) | Within locations | 84.62 | 83.22 | -1.40 |
| | Among provinces | 3.89 | 10.16 | +6.27 |
| | Among islands | 0.00 | 0.00 | +0.00 |
| | Among locations within province | 11.49 | 6.62 | -4.87 |

Values are the proportion (%) of total phenotypic variance attributable to each spatial level, estimated using linear mixed-effects models with Island, Province, and Location within Province as random effects (as in Table 6). Δ (pp) is the difference in percentage points (excluding East Nusa Tenggara minus full dataset).

**Supplementary Table S2. PERMANOVA results excluding East Nusa Tenggara.**

**Island-only model**

| Source | Df | Sum of squares | $R^2$ | F | P-value |
|---|---|---|---|---|---|
| Island | 9 | 475.89 | 0.153 | 10.222 | 0.001 |
| Residual | 510 | 2638.11 | 0.847 | – | – |
| Total | 519 | 3114.00 | 1.000 | – | – |

**Island + Province + Location model**

| Source | Df | Sum of squares | $R^2$ | F | P-value |
|---|---|---|---|---|---|
| Island | 9 | 475.89 | 0.153 | 10.830 | 0.001 |
| Province | 2 | 124.49 | 0.040 | 12.748 | 0.001 |
| Location | 4 | 52.76 | 0.017 | 2.701 | 0.002 |
| Residual | 504 | 2460.86 | 0.790 | – | – |
| Total | 519 | 3114.00 | 1.000 | – | – |

Results of permutational multivariate analysis of variance (PERMANOVA; 999 permutations) on standardized morphometric traits after exclusion of East Nusa Tenggara (n = 520).
